## Supplementary Materials for "Cell Marker Accordion: interpretable single-cell and spatial omics annotation in health and disease"

### SUPPLEMENTARY FIGURES

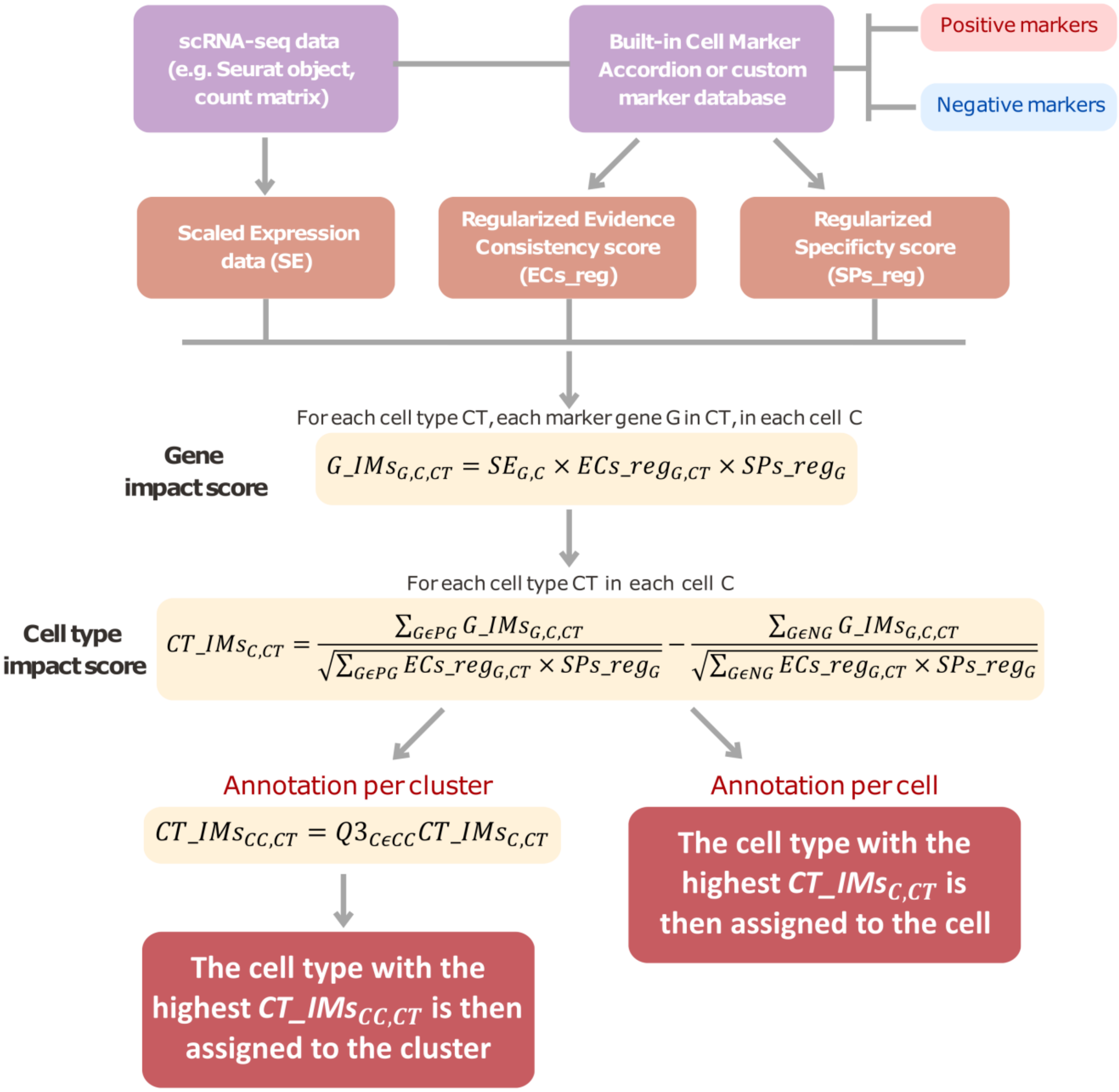


***Supplementary Fig.1: The Cell Marker Accordion: annotation and scoring workflow.*** *The figure outlines the Cell Marker Accordion annotation workflow: the necessary input, the main algorithmic steps and the primary annotation output (see* ***Methods****). SE: scaled expression data (e.g. Z-score); ECs: evidence consistency score; ECs_reg: regularized evidence consistency score; SPs: specificity score; SPs_reg: regularized specificity score; CT: cell type; G: marker gene; G_IMs: gene impact score, calculated for each cell type CT, each marker gene G, in each cell; PG: positive marker gene for cell type CT; NG: negative marker gene for cell type CT; CT_IMs: cell type impact score, calculated for each cell type CT in each cell; CC: cell cluster;* $CT\_IMs_{CC}$*: cell cluster cell type impact score.*

***Related to Fig.2***

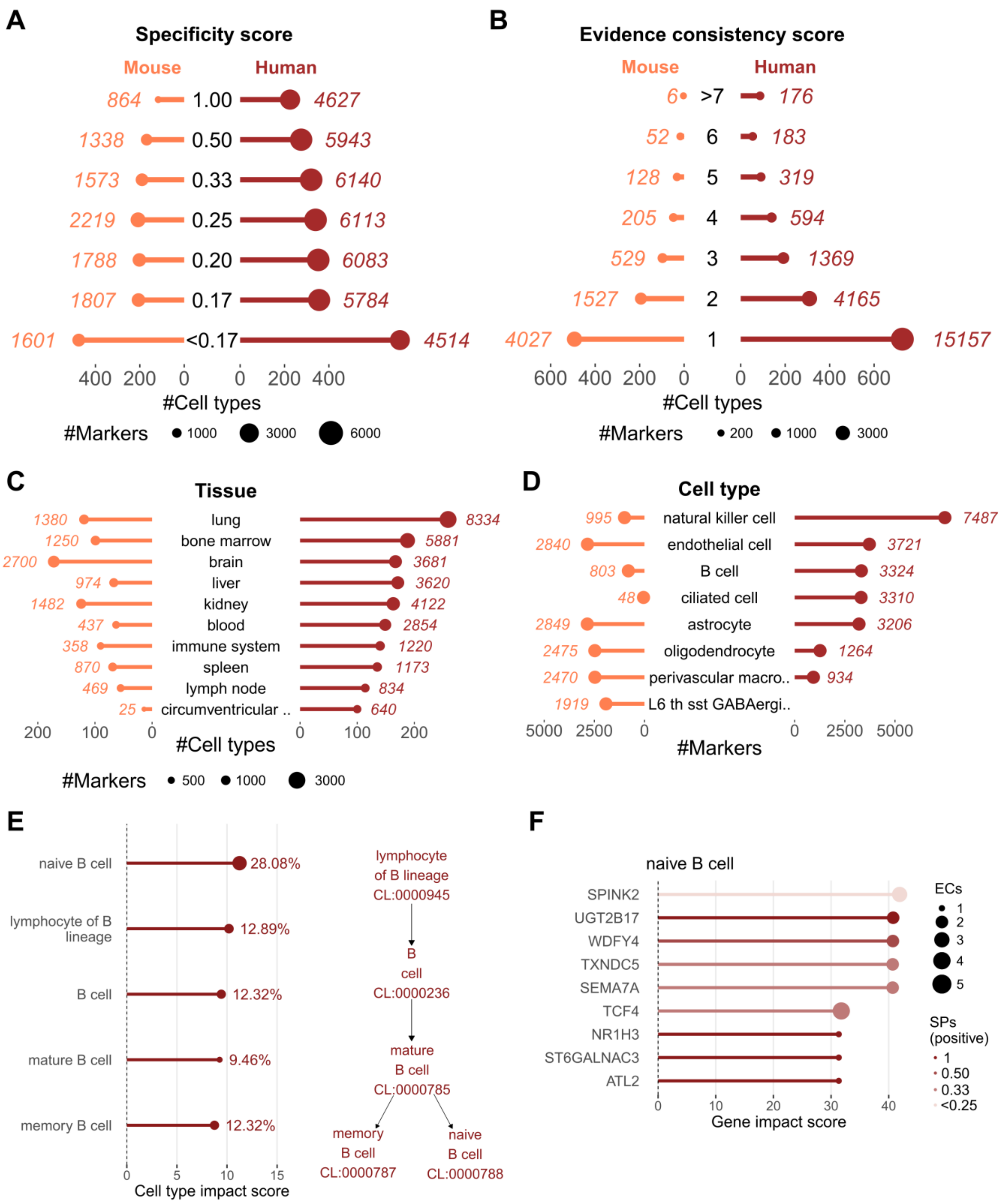


***Supplementary Fig. 2: The Cell Marker Accordion: scores and detailed annotation results. A*** *Number of cell types, reported on x axis, and number of markers, corresponding to dot size, for each specificity value in mouse (orange bars) and human (red bars).* ***B*** *Number of cell types, reported on x axis, and number of markers, corresponding to dot size, for each evidence consistency score value in mouse (orange bars) and human (red bars).* ***C*** *Cell Marker Accordion tissues with the highest number of markers for mouse (orange bars) and human (red bars).* ***D*** *Cell Marker Accordion cell types with the highest number of markers for mouse (orange bars) and human (red bars).* ***E*** *Example of cell types ranking output. On the left side, the top 5 cell types that compete for the annotation of*

*the same cluster are ordered according to their impact score. The percentage of cells labelled as the corresponding cell types is also shown. On the right side, the ontology tree of the top 5 cell types is represented.* ***F*** *Example of markers ranking output for a particular cluster. The most influential markers which drive the identification of the associated cell type, in this case, "naive B cell", are ordered according to their impact score. Dot size represents the evidence consistency score, and colors refer to marker specificity.*

***Related to Fig.2***

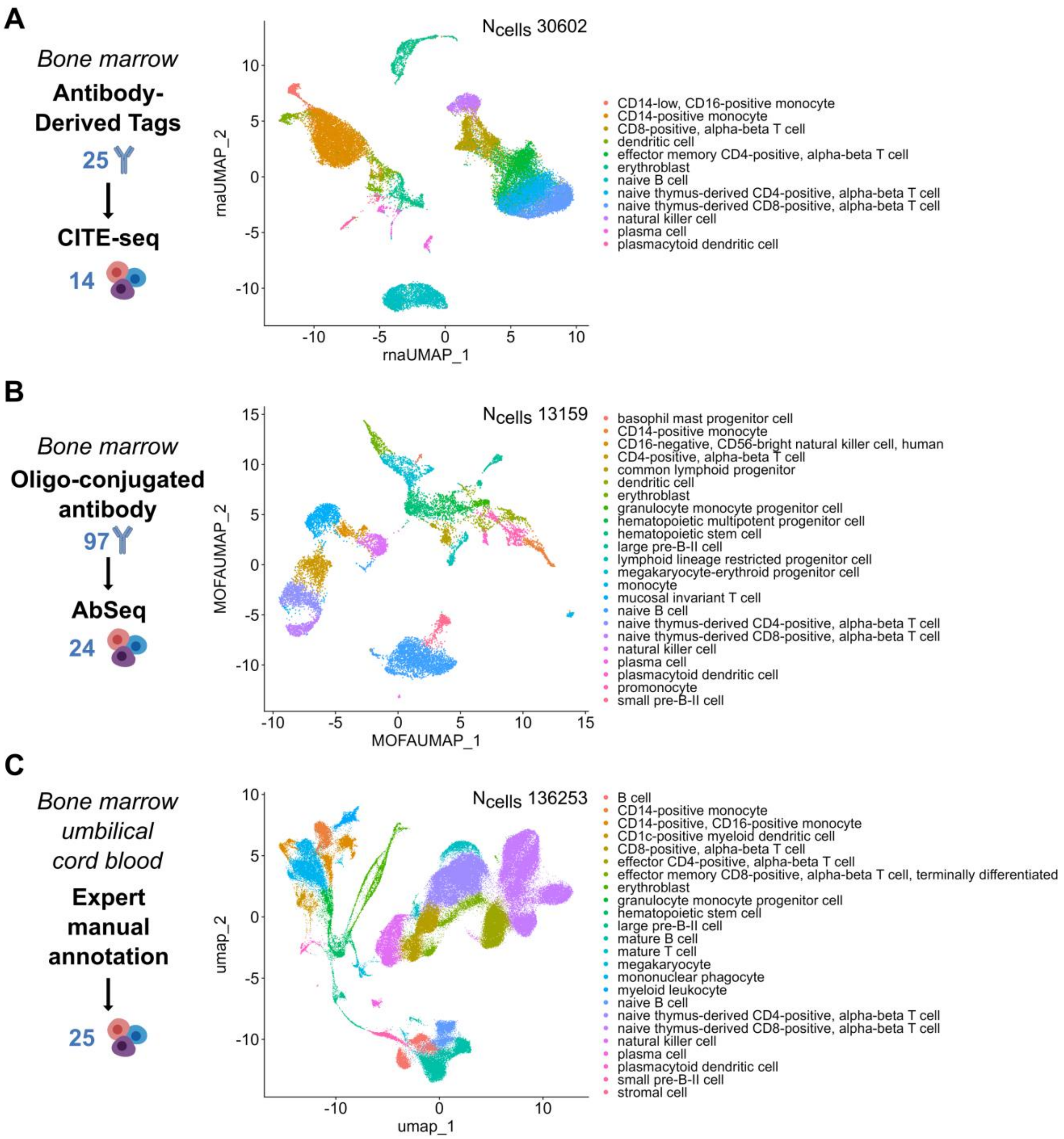


***Supplementary Fig.3: The Cell Marker Accordion annotation of cell types in complex single-cell multiomics.*** *Annotation with the Cell Marker Accordion of multi-omics single-cell datasets.* ***A*** *Human bone marrow dataset was obtained with a CITE-seq multi-modal approach (25 barcoded antibodies were used to quantify surface proteins and identify 14 different cell types, considered as the ground truth). Populations annotated by the Accordion are color-coded in the UMAP.* ***B*** *Human bone marrow dataset was obtained with an Ab-seq multi-modal approach (97 barcoded antibodies were used to quantify surface proteins and identify 24 different cell types, which is considered the ground truth). Populations annotated by the Accordion are color-coded in the UMAP.* ***C*** *Single-cell RNA-seq dataset of human cells from bone marrow and umbilical cord blood. Expert-based manual annotation identified 25 different cell types, considered as the ground truth.*

***Related to Fig.3***

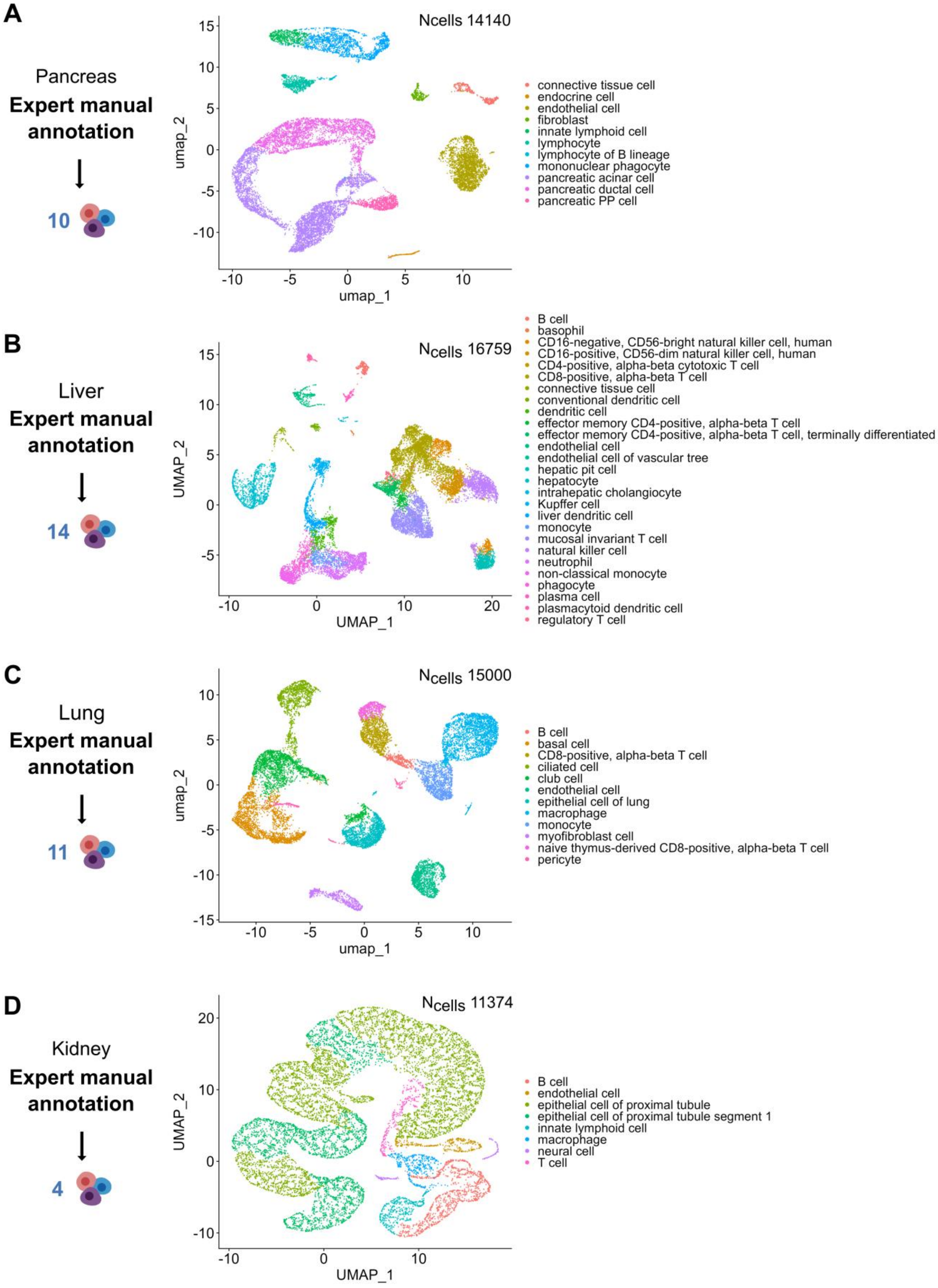


***Supplementary Fig.4: Cell Marker Accordion annotation of cell types in multiple tissues.*** *Annotation with the Cell Marker Accordion of single-cell datasets in multiple tissues* ***A*** *Single-cell*

*RNA-seq dataset of human cells from pancreas. Expert based manual annotation identified 10 different cell types, considered as the ground truth.* ***B*** *Single-cell RNA-seq dataset of human cells from liver. Expert based manual annotation identified 14 different cell types, considered as the ground truth. Populations annotated by the Accordion are color-coded in the UMAP.* ***C*** *Single-cell RNA-seq dataset of human cells from lung. Expert based manual annotation identified 11 different cell types, considered as the ground truth. Populations annotated by the Accordion are color-coded in the UMAP.* ***D*** *Single-cell RNA-seq dataset of human cells from kidney. Expert based manual annotation identified 4 different cell types, considered as the ground truth. Populations annotated by the Accordion are color-coded in the UMAP.*

***Related to Fig.3***

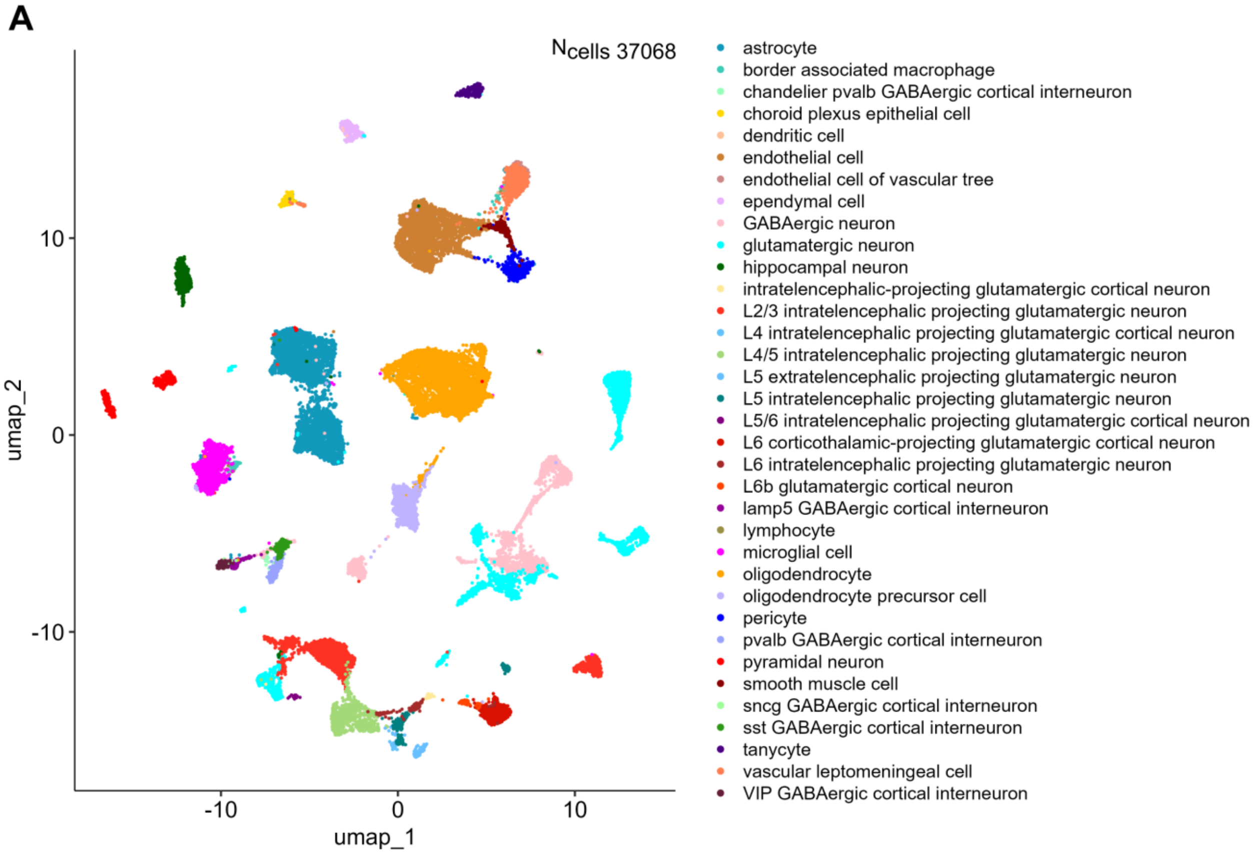


***Supplementary Fig.5: Mouse brain populations in spatial transcriptomic dataset. A** UMAP plot based on the transcriptional profile of each cell, with colors based on the original published annotation.*

***Related to Fig.4***

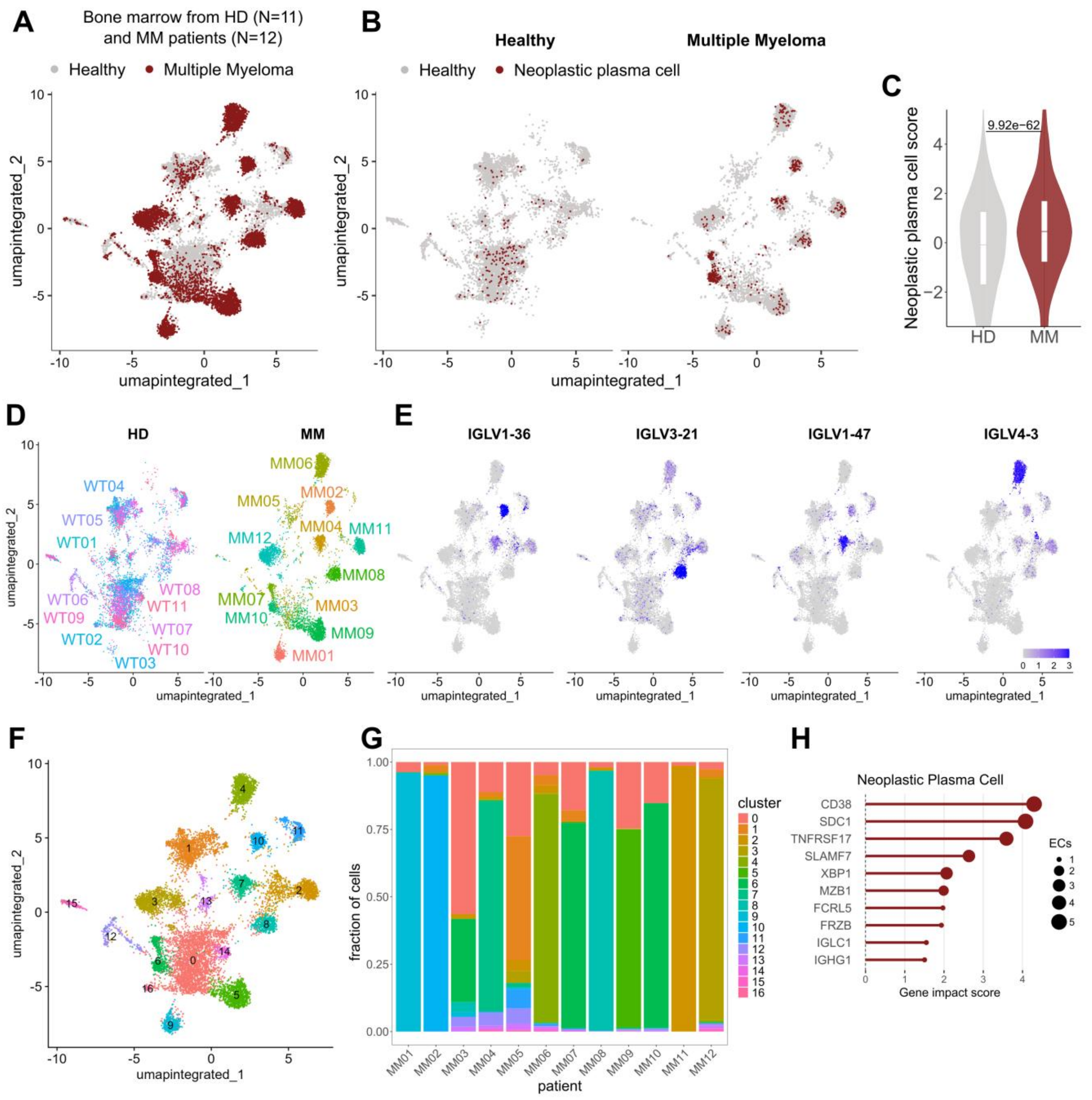


***Supplementary Fig. 6: The Cell Marker Accordion identifies myeloma plasma cells subtypes in multiple myeloma patients.*** ***A*** *Human bone marrow cells from healthy donors (HD) and multiple myeloma (MM) patients, for which bone marrow plasma cells were single-cell sorted by FACS and then sequenced.* ***B*** *Identification of neoplastic plasma cells with cell resolution. Cells are colored according to the neoplastic plasma cell score.* ***C*** *Distribution of neoplastic plasma cell scores. A significant increase is observed in MM patients.* ***D*** *UMAP showing HDs and MM patients cells. Cells from the same patients are color-coded.* ***E*** *Expression of Immunoglobulin variable region (IGVL) genes. Cells are colored according to gene expression levels.* ***F*** *UMAP visualization of cells of HD and MM patients, with clusters color-coded and labelled.* ***G*** *Cluster composition in each MM patient.* ***H*** *Marker genes with the highest impact in defining neoplastic plasma cells from MM patients. One-tailed Wilcoxon Rank Sum test was used for panel* ***C****. P-value is displayed.*

***Related to Fig.5***

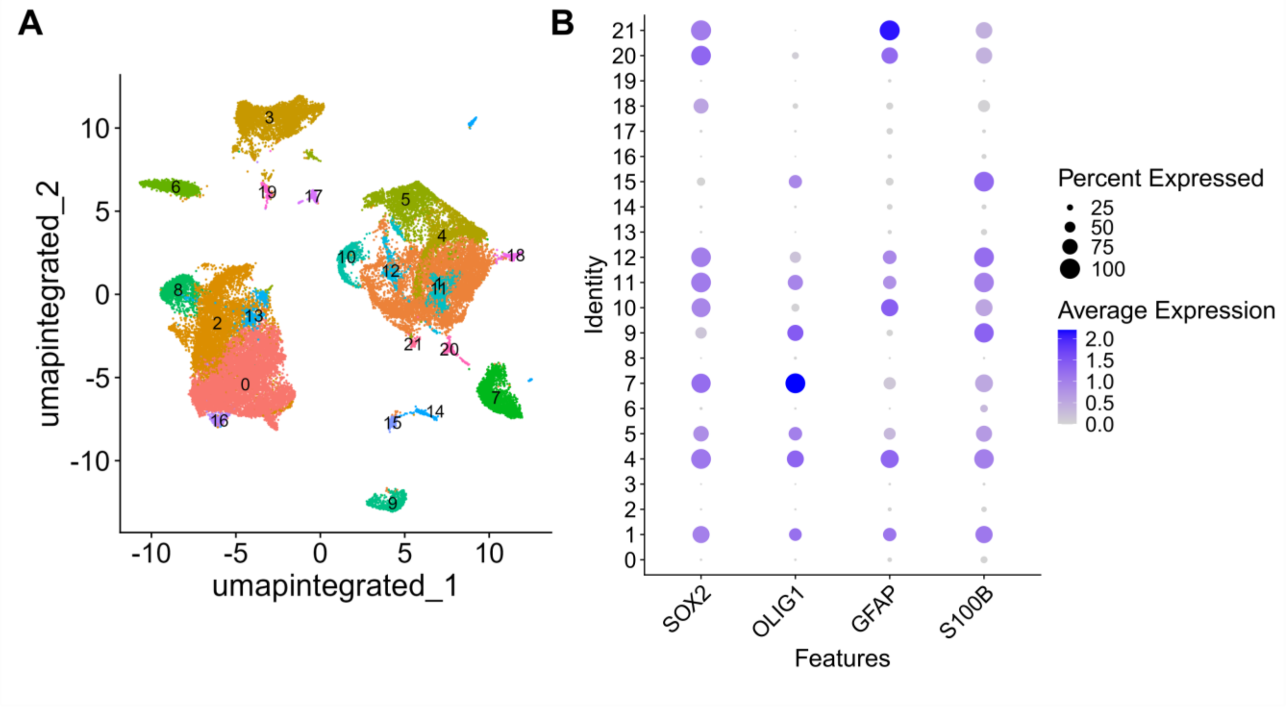


***Supplementary Fig.7: Clusters of tumoral cells in glioblastoma patients. A*** *UMAP visualization of cell clustering from glioblastoma patient samples, with clusters color-coded and labelled.* ***B*** *Expression levels of tumor markers across the identified clusters.*

***Related to Fig.6***

### SUPPLEMENTARY DATA

**Supplementary Data 1.** List of available gene marker databases, resources and integration procedure used to build the Cell Marker Accordion

**Supplementary Data 2.** Comparison of available marker-based annotation tools in terms of features and capabilities.

**Supplementary Data 3.** Cell type specific performance annotation of single-cell and multi-omic datasets.

**Supplementary Data 4.** List of all published datasets used in this study

**Supplementary Data 5.** Tools and parameters used for benchmarking analysis

**Supplementary Data 6.** MDS patient characteristics and sequencing metrics

**Supplementary Data 7.** Innate immune response gene signatures

**Supplementary Data 8**. The Cell Marker Accordion database

### SUPPLEMENTARY METHODS

#### Single-cell data analysis

All single-cell datasets were analyzed employing a standard pre-processing pipeline for single-cell RNA-seq data, using Seurat (version 5.0.3) in the R environment (version 4.2.3). Data were first log-normalized and the most highly variable features were identified and scaled. Linear dimensionality reduction through Principal Component Analysis (PCA), was performed on scaled data and scRNA-seq cells were clustered based on a shared nearest neighbor graph (obtained with the FindNeighbors function) employing the FindClusters function with the default Louvain algorithm for community detection. For cluster visualization Uniform Manifold Approximation and Projection for Dimensionality Reduction (UMAP) was employed. In case of scRNA-seq datasets integration, the IntegrateLayers function was used to align cells across multiple samples by correcting batch effects while preserving biological variability. The MERFISH mouse brain dataset was analyzed employing a standard pre-processing pipeline for imaging-based spatial dataset, using Seurat (version 5.0.3) in the R environment (version 4.2.3). Data were first normalized with SCT-transform. Linear dimensionality reduction through Principal Component Analysis (PCA), was performed on transformed data and cells were clustered based on a shared nearest neighbor graph (obtained with the FindNeighbors function) employing the FindClusters function with the default Louvain algorithm for community detection. For cluster visualization Uniform Manifold Approximation and Projection for Dimensionality Reduction (UMAP) was employed. The mouse coronal brain section with ID "Zhuang-ABCA-1.080" was selected based on the highest number of barcodes (37,068). Spatial mapping of cells was performed using NIfTI images, aligning the spatial coordinates of the cells to the Allen CCF 2020 with a voxel spacing of 0.01 mm.
